# Parallel evolution of phage resistance - virulence trade - offs during *in vitro* and nasal *Pseudomonas aeruginosa* phage treatment

**DOI:** 10.1101/2021.09.06.459069

**Authors:** Meaghan Castledine, Daniel Padfield, Pawel Sierocinski, Jesica Soria Pascual, Adam Hughes, Lotta Mäkinen, Ville-Petri Friman, Jean-Paul Pirnay, Maya Merabishvili, Daniel De Vos, Angus Buckling

**Affiliations:** College of Life and Environmental Sciences, Environment and Sustainability Institute, University of Exeter, Penryn, Cornwall, TR10 9FE, U.K.; Department of Biology, University of York, York, UK; Laboratory for Molecular and Cellular Technology, Queen Astrid Military Hospital, Brussels, Belgium

## Abstract

With rising antibiotic resistance, there has been increasing interest in treating pathogenic bacteria with bacteriophages (phage therapy). One limitation of phage therapy is the ease at which bacteria can evolve resistance. Negative effects of resistance may be mitigated when resistance results in reduced bacterial growth and virulence, or when phage coevolve to overcome resistance. Resistance evolution and its consequences are contingent on the bacteria-phage combination and their environmental context, making therapeutic outcomes hard to predict. One solution might be to conduct “*in vitro* evolutionary simulations” using bacteria-phage combinations from the therapeutic context. Overall, our aim was to investigate parallels between *in vitro* experiments and *in vivo* dynamics in a human participant. Evolutionary dynamics were similar, with high levels of resistance evolving quickly with limited evidence of phage evolution. Resistant bacteria – evolved *in vitro* and *in vivo* - had lower virulence. *In vivo*, this was linked to lower growth rates of resistant isolates, whereas *in vitro* phage resistant isolates evolved greater biofilm production. Population sequencing suggests resistance resulted from selection on *de novo* mutations rather than sorting of existing variants. These results highlight the speed at which phage resistance can evolve *in vivo*, and *in vitro* experiments may give useful insights for clinical evolutionary outcomes.

## Introduction

By 2050, as many as 10 million people may die annually from antimicrobial resistant infections (WHO, 2019). To combat this threat, phage therapy is being increasingly researched as a complement to antibiotics (Abedon, 2019; Kwiatek et al., 2020). Phage therapy uses viruses (phage) that infect pathogenic bacteria to clear infections (Kwiatek et al., 2020). Phage therapy has several advantages to antibiotics by being species specific (thus not harming beneficial microbiota) (Koskella and Meaden, 2013) and self-amplifying at infection sites (Górski et al., 2015). A major consideration with the use of therapeutic phages is the ease and speed (sometimes within hours) at which bacteria can evolve resistance, which can greatly limit the therapeutic potential of phages (Jariah and Hakim, 2019; Torres-Barceló, 2018; Torres-Barceló and Hochberg, 2016). However, phages may contribute to reducing the severity of infections even after resistance evolves. First, phage can sometimes evolve to overcome this resistance (i.e. bacteria-phage coevolution) and so they may continue lowering bacterial densities (Gandon et al., 2008; Hampton et al., 2020). Second, resistance can trade-off against bacterial growth rates and virulence; as the alteration or loss of bacterial surface structures, typically associated with phage resistance, can affect other fitness-related phenotypes such as nutrient uptake, biofilm formation and binding of host receptors (Mangalea and Duerkop, 2020). Predicting the evolutionary outcome of phage therapy is therefore of crucial importance (Abedon, 2017).

Bacteria-phage dynamics and its consequences for phage therapy are highly context-specific (Gandon et al., 2008; Hampton et al., 2020). Both human clinical and animal models studies report a range of outcomes with differing phage therapy success, and differences in the extent of resistance evolution and associated costs and virulence (Abedon, 2019; Brix et al., 2020; Kwiatek et al., 2020; Leitner et al., 2021). One solution to better understand the outcome of *in vivo* therapy is to conduct controlled *in vitro* experiments with the relevant organisms. However, while most of our understanding of the ecology and evolution of bacteria-phage interactions comes from highly controlled *in vitro* environments (Gandon et al., 2008; Hampton et al., 2020), we do not know how well *in vitro* dynamics mirror *in vivo* dynamics quantitatively or even qualitatively.

There are good reasons to expect the outcomes may be very different. *In vivo* environments are expected to be more stressful for bacteria owing to limiting resources, presence of immune cells and competing microbiota (Everest, 2007). Consequently, these conditions may differ significantly from studies of bacteria-phage coevolution in nutrient rich microcosms which could drive differences in (co)evolutionary dynamics and associated trade-offs (Gandon et al., 2008; Hampton et al., 2020). In order to evolve phage resistance, bacteria must have sufficient mutation rates supplied either by high resources (facilitating rapid growth) or population mixing (genetic exchange) (Gandon et al., 2008; Hampton et al., 2020; Morgan et al., 2010, 2005; Pal et al., 2007). Smaller population sizes *in vivo* may therefore limit resistance. Moreover, resistance costs and trade-offs may be greater as a consequence of the additional selection pressures facing bacteria, which may limit selection for resistance, making it easier for phage to overcome resistance, or reduce growth rates and virulence (Buckling et al., 2006; Scanlan et al., 2015).

Consequently, the main aim of this research was to determine how well *in vitro* bacteria – phage coevolutionary dynamics mirror dynamics in a clinical context. As a starting point, we followed bacteria - phage coevolutionary dynamics during a phage-mediated decolonisation trial in parallel with the dynamics between the same bacteria and phage *in vitro*. The nasal cavity can act as a source of infection, notably from highly drug-resistant *Pseudomonas aeruginosa* and *Staphylococcus aureus* when patients undergo hospitalisation and are treated with broad spectrum antibiotics (Mainz et al., 2009; Wertheim et al., 2004). The trial was conducted to try and decolonise intensive care (ICU) patients with *P. aeruginosa* and *S. aureus*, using a phage cocktail prior to antibiotic treatment in prevention of ICU infections with these pathogens. Mucirocin (trade name Bactroban) and chlorhexidine are routinely used for nasal decolonisation in such contexts, but extensive use of these agents has led to resistance and new nasal antiseptics need to be considered (Edgeworth, 2011). We focused on a single patient, in which we could detect *P. aeruginosa* at the beginning of the trial and both *P. aeruginosa* and phage during treatment. Decolonisation of *P. aeruginosa* was ultimately successful in this patient. We used the pre-treatment *P. aeruginosa* isolates and evolved these populations *in vitro* in the absence and presence of treatment phages. We measured bacteria-phage (co)evolution and while we found extensive resistance evolution both *in vivo* and *in vitro*, we only found evidence of limited phage infectivity evolution *in vitro*. We determined the consequences of resistance, with respect to virulence, and the possible mechanisms linking this resistance – virulence trade – off using phenotypic (growth rate, biofilm assays) and metagenomic analyses for the *in vivo* and *in vitro* populations. As we only had access to samples from a single patient, we did not have a control group to understand the impact of phage application frequency *in vivo*. However, we were able to do this in our *in vitro* experiments, where we included two phage treatments: one where phages were added repeatedly (mimicking multiple phage doses during *in vivo* phage treatment) and one where phage were added once (at the start of the experiment). By comparing these treatments and to the *in vivo* treatment, we were able to gain further insight into how multiple phage doses may affect phage therapy.

## Methods

### Ethics statement

The decolonisation study protocol was approved by the Leading Ethics Committee of the “Université Catholique de Louvain” (Avis N°: B-403201111110). The study was performed in accordance with the ethical standards as laid down in the Declaration of Helsinki and as revised in 2013. The patients gave informed consent and their anonymity was preserved.

### Patient information, bacteria and phage strains and phage-bacteria evolution *in vivo*

Bacterial isolates were sampled from a patient as part of a clinical trial in Brussels, Belgium. The patient (85 years old, male) had been admitted to an intensive care unit in 2015 for treatment of burns (10% body area). Independent of his burns, a nasal colonisation of *P. aeruginosa* was observed. Here, the patient was treated every day with a three-phage cocktail (BFC-1) for five days, which consisted of two phages (14-1 and PNM) that could lytically infect *P. aeruginosa*. The phage cocktail included a third phage that specifically infects *S. aureus* bacterium but was not detected in this patient. This phage cocktail has been characterised to ensure no prophage elements are present or that the phage carry toxin-encoding genes that may increase bacterial resistance and/or virulence (Merabishvili et al., 2009). Furthermore, a purification process including the removal of endotoxins in phage production was used to ensure such waste products cannot harm patients (Merabishvili et al., 2009). As 14-1 infects via a lipopolysaccharide (LPS) (Betts et al., 2014; Kropinski et al., 1977) receptor and PNM uses type IV pili (Ceyssens et al., 2011), resistance to one phage is unlikely to result in cross-resistance and neither phage should compete for receptor sites. Swabs of both nasal cavities were taken on day 0 (commencement of phage therapy), 2, 4, 7 and 13 and frozen at -80 °C in lysogeny broth containing 20% glycerol. Individual *P. aeruginosa* isolates were obtained by plating swabs onto cetrimide selection plates. Colonies were picked and inoculated into Luria broth (LB) and grown overnight, shaking at 180 r.p.m., at 37 °C to achieve high cell densities. Additionally, swabs were soaked in LB broth to amplify phage overnight in the same conditions. Phage was isolated from overnight cultures via chloroform extraction: 100 µL of chloroform was added 900 µL of culture, vortexed and then centrifuged (14000 r.p.m., 1 min); the supernatant was isolated and stored at 4 °C. Culture samples were frozen in 80% glycerol (final concentration 40%) at -80 °C. Spot assays on lawns of *P. aeruginosa* P573 (a phage susceptible control strain) confirmed the presence of phage at all time-points. Twelve individual plaques were picked at each time-point and amplified with an overnight culture of P573 to obtain clonal isolates. Individual isolates of *P. aeruginosa* were isolated from days 0 (n = 24), 2 (n = 8) and 4 (n = 2) for use in this study and swabs were negative on days 7 and 13. Individual isolates were treated as biological replicates for *in vivo* study.

### Experimental *in vitro* phage-bacteria evolution

To examine whether findings *in vitro* are consistent with *in vivo* bacteria-phage (co)evolution, phage therapy was replicated in the laboratory. We used two treatments: (1) phage added “once”, at the start of the experiment, and (2) phage added at every second transfer (“repeated” treatment). The repeated phage treatment aimed to emulate the conditions of the clinical trial while the “once” condition was used to assessed whether repeated doses change the observed dynamics. The phages were identical to the two lytic phages (14-1 and PNM) used in the clinical trial. A control group in which bacteria evolved in the absence of phage was set up and there were 6 biological replicates in the control and each phage treatment.

Prior to experimental set-up, *in vivo* ancestral isolates (n = 24, same isolates as day 0) were individually amplified in LB media and cultured at room temperature for two days. Cell densities were estimated via optical density (wavelength 600 nm; OD_600_) measures. Cultures were centrifuged (14000 r.p.m., 3 min), the supernatant removed, and resuspended in M9 buffer to equalise cell densities. These were added to a master-mix in which isolates were equally represented in equal ratio (total 10^8^ CFUs / mL; ∼4 × 10^6^ CFUs / mL of each isolate). 100 µL of master-mix (10^7^ CFUs; ∼4 × 10^5^ CFUs per isolate) was inoculated into 5 mL LB media. 100 µL of phage was added to treatment microcosms at a total multiplicity of infection (MOI) of 0.5. Microcosms were cultured, shaken (180 r.p.m.), at 37 °C. 1% transfers (50 µL culture into 5 mL LB) took place every two days for a total of 12 days. At each transfer, culture samples were frozen in glycerol, and in the phage treatments, phage were extracted as described previously. Cultures were diluted and plated from frozen stocks onto LB agar and incubated for two days at 37 °C. From each biological treatment replicate and time-point (days 4, 8 and 12), 16 bacterial colonies were picked (96 total colonies) for time-shift assays to measure coevolutionary changes. An additional 24 isolates were isolated at day 12 from each treatment and control group replicate for bacterial virulence assays. Individual isolates are treated throughout as biological replicates nested within their treatment (biological) replicate. Picked isolates were individually amplified in LB media overnight before being frozen as described previously.

### Measuring resistance-infectivity (co)evolution

To assess whether bacteria and phage had coevolved *in vivo* and *in vitro*, resistance assays were conducted where bacteria were exposed to phage populations from the same/contemporary, past, and future time-points (time-shift assays) (Buckling and Rainey, 2002). If bacteria and phage coevolved, both bacteria and phage would show significant changes in resistance and infectivity, respectively, through time. Increases in resistance and infectivity would be indicative of arms race dynamics (ARD), while non-directional changes would be indicative of fluctuating selection dynamics (FSD) (Lopez Pascua et al., 2014). Infectivity/resistance was determined by spot assays in which 100 µL of overnight bacterial cultures was added to 5-8 µL soft LB agar overlay and 5 µL of each ancestral phage, or evolved phage mixture, was spotted onto the overlay. For *in vivo* isolates, ancestral isolates (isolated pre-phage therapy) and bacterial isolates from days 2 and 4 were tested against the phage clonal isolates from days 2, 4 and 7. A bacterial isolate was scored as susceptible (1 = susceptible, 0 = resistant) if at least one of twelve clonal phage isolates could infect. *In vitro* isolates from days 4, 8 and 12 were tested (16 colonies per time-point) against phage populations isolated from each time-point. In addition, we assessed the infectivity of both ancestral phage against the ancestral bacteria (n = 24), the *in vivo* isolates (n = 8), and the *in vitro* populations from day 12 (n = 24 per treatment replicate).

### Virulence assays

Changes in bacterial virulence were measured using a *Galleria mellonella* infection model (Tsai et al., 2016). Virulence of all ancestral (n = 24) and *in vivo* isolates (n = 10) were measured, however two *in vivo* isolates (from day 2 of phage therapy) were excluded from analyses as time of death was not accurately determined. As few isolates remained phage susceptible (n = 3), these were pooled with phage resistant isolates to determine a change in virulence from pre-(ancestral, n = 24) to post-phage treatment (n = 8). For *in vitro* isolates, where possible, one resistant and one susceptible (to ancestral phage) isolate were selected from each treatment replicate to give a total of 12 resistant and 12 susceptible isolates, with 14 isolates from the “phage added once” and 10 isolates from the “phage added repeatedly”. As some treatment replicates contained no susceptible isolates, multiple resistant isolates were chosen (Table S1). Seven *in vitro* isolates evolved resistance to one phage only (single resistance), therefore their virulence was measured to assess whether changes in virulence were mediated by resistance to one or both phages. The virulence of *in vitro* isolates was contrasted to six isolates from the control group isolated from 6 independent replicates.

For each bacterial isolate (total of 56 biological replicates), 20 *G. mellonella* larvae (20 technical replicates) were infected with 10 µL (∼100 CFUs) of overnight culture (cells resuspended in M9 buffer) using a sterile syringe. Twenty *G. mellonella* larvae were injected with 10 µL of M9 buffer as a technical control. Survival of each larvae was assessed every two hours for up to 58 hours.

### Growth rate assays

We measured bacterial growth rates as a potential mechanism underlying virulence variation and cost of resistance for ancestral and all evolved isolates (*in vitro* and *in vivo*) tested in the virulence assay. Growth curves were measured in a 96 well plate with 10 µL of culture (1.0 × 10^5^ CFUs) from each isolate inoculated into individual wells containing 140 µL LB broth. 26 wells remained as blank controls. The plate was incubated in an OD reader (Biotek Instruments Ltd) at 37 °C in shaken conditions and OD_600_ was measured every 20 mins for 24 hours to estimate changes in cellular density. Growth curves were independently replicated 3 times (technical replicates).

### Biofilm assay

Biofilm production of isolates isolated for virulence and growth rate assays was measured using a resazurin assay, based upon a previously described protocol (Peeters et al., 2008). Overnight monocultures were diluted to 0.1 OD_600_ (∼10^9^ CFU/mL) into M9 buffer. 100 µL of each culture was inoculated into a sterile 96 well plate and incubated at 37 °C for 4 hrs for cellular adhesion. Then, the supernatant was removed, and wells were washed gently with 100 µL of M9 buffer by pipetting up and down 3 times. 100 µL of fresh LB media was added to each well including 25 controls (no bacteria) and plates were incubated for 16 hrs. Liquid was removed and wells were washed with 100 µL M9 buffer. 100 µL of fresh M9 was added followed by 10 µL resazurin solution (final resazurin concentration: 100 µM). Fluorescence (*λ*ex: 560 nm and *λ*em: 590 nm) was measured after 1 hr incubation. Biofilm assays were independently replicated 3 times (technical replicates).

### Sequencing

We sequenced the genomes of the bacterial isolates used for the *in vitro* experiment to establish to what extent evolution was driven by *de novo* mutation or selection on existing genetic variation. This is especially pertinent as the experiment did not start from a clonal population, rather from 24 isolates (which included an isolate that was resistant to one phage). At the end of the experiment, we picked 24 isolates, extracted DNA from isolates individually and performed whole genome sequencing (WGS) on pools of the bacterial isolates that were isolated from each replicate (pool-seq). Total DNA extraction (19 pooled-isolate samples) was performed using the “Qiagen QIAmp DNA Mini Kit” following the manufacturer’s instructions. An Illumina HiSeq 2000 sequencer was used to generate 100 bp paired reads from a 500 bp insert library.

Reads were trimmed for the presence of Illumina adapter sequences and for quality scores below 10 using *TrimGalore* (v0.6.4_dev, powered by *Cutadapt*). *TrimGalore* also removed any paired end reads when one of the paired end reads was shorter than 20 bp. Trimmed reads were mapped to the *P. aeruginosa* UCBPP-14 reference genome (Locus: NC_008463) with *bwa-mem* (v0.7.17-r1188), and duplicates were marked using *samblaster* (v0.1.24). Subsequent bam files were split into mapped and unmapped reads, sorted and indexed using *samtools* (v0.1.19-44428cd). Across all files, ∼93% of the reads mapped to the reference genome (minimum 92.5%), giving on average ∼12,000,000 reads per replicate (minimum 8,807,569). Variants were called identified using *freebayes* (v1.3.2-dirty) with ploidy set to 24 (*-p 24*) and we assumed pooled samples with a known number of genome copies (*--pooled-discrete*). Due to the high ploidy of the pooled samples, we reduced the number of alleles that are considered at the cost of sensitivity to lower frequency alleles at multiallelic loci by setting *--use-best-n-alleles 4*.

Next, we compiled the vcf files (standard output of SNP callers) into *R* using the package *vcfR* and kept only the variants which were sequenced at a depth within two standard deviations of the mean. Variants were further filtered to retain only those with quality scores >30. We used the two replicate sequencing runs of the same samples to conservatively identify variants; variants were only kept if they occurred in both sequencing runs and if there was less than 0.3 difference in frequency between runs. We then only kept variants that had different frequency than the ancestor (> 1/24 difference in frequency). To identify significant differences in genetic variants (SNPs and indels) between treatments, we first filtered unique that were present in at least three replicates of any treatment. This identified 960 potential variants. We then performed a Wilcoxon test on each variant, with frequency as the response variable and treatment (control, phage added once, and phage added repeatedly) as the predictor. Significant genetic variants were identified if the adjusted p value (*fdr* method) was < 0.05. Of these significant changes, 160 were existing genetic variants and 120 were novel mutations. We then checked where each variant was in the genome and only retained variants that were occurred in a gene. After these steps, 50 genetic variants were identified in genes of known function, and 236 in genes of unknown function.

For the *in vivo* isolates, DNA extraction and whole genome sequencing on each isolate individually by MicrobesNG using in-house protocols (“MicrobesNG Sequencing Service Methods v20210419,” n.d.). Whole genome sequencing was carried out on each isolate individually by MicrobesNG using Illumina sequencers (HiSeq/NovaSeq) to create 250 bp paired end reads. In-house processing by MicobesNG trimmed the adapters and removed reads with a quality < 15. We then processed the reads as for the pooled sequencing, except ploidy was set to 1 when using *freebayes*. On average, 81% of reads mapped back to the reference genome, but there was more variation as compared to the *in vitro* experiment, with a minimum of 44% and a maximum of 94%. As we only had isolates from a single patient, there was limited statistical analysis we could do, so we primarily concentrated on whether the significant changes found *in vitro* were also present in the *in vivo* isolates.

### Statistical analyses

All data was analysed using R (v. 4.0.3) in RStudio (Team, 2013) and all plots were made using the package “*ggplot2*” (Wickham, 2016). Model simplification was conducted using likelihood ratio tests and Tukey’s post-hoc multiple comparison tests were used to identify the most parsimonious model using the R package “*emmeans*” (Lenth, 2018). First, we established whether bacteria and phage had coevolved *in vivo* and in each *in vitro* treatment using general linear mixed effects models. Models were fitted using the package “*lme4*” (Bates et al., 2014). *In vivo* isolates from day 4 were excluded as their small sample size (n = 2) prevented model convergence. For *in vivo* isolates: infection (1 = susceptible, 0 = resistant) was analysed against phage time-point (fixed effect) with a binomial error structure. A random effect of “isolate” was included to account for the same bacterial isolate being used in infectivity assays across each phage time-point. For *in vitro* analyses, the proportion of isolates infected was analysed against fixed effects of phage and bacterial time-points with separate models for each phage treatment. To see if there were any differences in phage resistance to ancestral phage or *in vitro* phage isolates at the end of the experiment, the proportion of isolates infected was analysed against fixed effects of phage treatment and phage type (ancestral or end – point phage). All *in vitro* models included a random effect of “replicate” (which treatment replicate isolates had been isolated from) with a binomial error structure.

Virulence of bacterial isolates was analysed using survival curves from the survival of *Galleria* infected with different isolated isolates. Survival curves were fitted using Bayesian regression models using the package “*rstanarm*” (Goodrich, 2020) and parameters were estimated using “*tidybayes*” (Kay and Mastny, 2020). For all models, we fit a proportional hazards model with an M-splines baseline hazard. Models were ran for 3000 iterations and 3 chains were used with uninformative priors. Model convergence was assessed using Rhat values (all values were 1) and manually checking chain mixing. For all models, log hazards were estimated for each treatment value and hazard ratios were calculated as the exponential of the difference between two log hazards. A hazard ratio below one indicates a decrease in virulence compared to the baseline treatment, with a value above one indicating an increase in virulence. Median hazard ratios with 95% credible intervals that do not cross 1 indicate a significant difference in virulence between two factors. Proportion of *Galleria* that died and mean time to death of *Galleria* that died were calculated as summary statistics for each treatment.

For virulence analyses of *in vivo* isolates, treatment (pre- or post-phage therapy) was the only predictor of survival, with a random effect of “isolate” added to account for multiple *G. mellonella* larvae being infected by the same isolate. For the *in vitro* isolates, survival was analysed against interacting fixed effects of resistance (resistant, single resistant, susceptible) and treatment (no phage added, phage added once, phage added repeatedly). Here, an additional random effect of “replicate” was included to account for clonal non-independence.

Biofilm productivity of *in vitro* isolates was estimated using linear mixed effects models. Biofilm production (fluorescence 560/590nm) was log transformed to account for the data being abundance-based. Biofilm production was averaged over each technical replicate for each biological replicate (individual bacterial isolates). Biofilm production was used as the response variable, and explained with interacting fixed effects of treatment and phage resistance, while treatment replicate was included as a random effect. One outlier was removed each from the susceptible and control groups as these were likely contamination (full data cleaning process and impact on parameter estimates explained in the SI and effects of outlier removal on estimates shown in Tables S3 & S4). Due to the low number of isolates, biofilm productivity of *in vivo* isolates from before and after phage treatment as well as phage susceptible and resistant isolates were estimated using a non-parametric bootstrap using the R package “*boot*” (Canty and Ripley, 2021). Non-overlapping 95% CIs were interpreted as being significantly different.

Before growth rate analyses, data points from the first 2.5 hrs were removed as a sharp increase in OD was observed before lag phase independent of exponential growth (Figure S1). Growth rates were estimated using a rolling regression taking the steepest slope of the linear regression between *ln* OD_600_ and time in hours in a shifting window of every 7 time points (every 2.3 hrs). Growth rate was averaged over each technical replicate for each biological replicate (individual bacterial isolates). For *in vitro* isolates, growth rate was analysed against treatment and resistance in a linear mixed effects model with a random effect of treatment “replicate”. Growth rates of isolates pre- and post-phage treatment, and resistant and susceptible *in vivo* isolates were compared using 95% confidence intervals (CIs) that were estimated using a non-parametric bootstrap.

To look at genetic differences *in vitro*, we calculated several commonly used metrics using the frequency data of the 50 variants in known genes that differed significantly between our treatments: (1) the genetic distance from the ancestral population, calculated as the sum of the difference of the proportion of each SNP/indel in each population from the ancestral proportion; (2) the number of unique *de novo* SNPs/indels in each population; (3) alpha diversity, calculated using a modified version of the Hardy-Weinberg equilibrium, such that 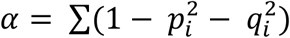, where *i* is the position of each SNP/indel, *p* is the proportion of the SNP/indel and *q* is 1 – *p*. This is equivalent to expected heterozygosity. Differences between these metrics were analysed using linear models. To test whether consistent genetic differences occur within treatments, we performed non-metric multidimensional scaling on the Euclidean distance matrix of SNPs/indels and their proportions in each population using “*metaMDS*” in the R package “*vegan*”. Nonmetric multidimensional scaling aims to collapse information from multiple dimensions (*i*.*e*., from different populations and multiple SNPs/indels per population) into just a few, allowing differences between samples to be visualized and interpreted. Permutational ANOVA tests were run using “*vegan::adonis*”, with Euclidean distance as the response term and treatment as the predictor variable with 9999 iterations. Changes in beta-diversity were examined using “*vegan::betadisper*” using the same response and predictor variables as in the PERMANOVA.

For the *in vivo* sequencing, we looked for the presence of variants found to be significant *in vitro* in the *in vivo* isolates. To see if any of the genetic changes could explain differences in phenotypes of the isolates, we did non-metric multidimensional scaling as described previously. Phenotypic traits of the isolates (virulence, growth rate, resistance profile, and biofilm production) were then mapped onto the resulting NMDS plot to see whether observed genetic changes correlated with any changes in phenotype (Figure S2).

## Results

### Resistance and infectivity (co)evolution

#### In vivo

Resistance to phage can evolve via *de novo* mutations or selection on pre-existing resistance and we therefore tested the resistance of ancestral bacterial isolates to ancestral phage and pooled phage isolates isolated *in vivo*. Of the 24 ancestral isolates isolated *in vivo* at day 0, 23 were susceptible to both the ancestral phage (14-1 and PNM) while one was resistant to 14-1 but susceptible to PNM. In subsequent tests of phage resistance to phage isolates isolated *in vivo*, all ancestral bacteria were susceptible to phage (Figure 1a). We considered whether bacteria and phage coevolved *in vivo* using time-shift assays to determine changes in resistance and infectivity through time. After two days, we saw evidence of resistance evolution, with more than half of bacterial isolates being resistant to ancestral phage. 1 of 2 isolates isolated at day 4 were phage resistant and no *P. aeruginosa* was detected from nasal swabs after this time point. Although both bacterial populations from days 2 and 7 showed observationally higher phage resistance (87.5% day 2, 100% day 4) against day 2 phage, this was non-significantly different to resistance shown to ancestral phage and phage from days 4 and 7 (Tukey HSD: p > 0.05; Table S2; Figure 1a). This non-significance is likely due to low statistical power from limited clonal numbers (only 8 isolates from day 2) and the consistency of resistance levels in bacterial populations from day 2 apart from to its contemporary phage; overall, results showed equal levels of phage resistance over time *in vivo*. The absence of changes in phage infectivity through time did not warrant further investigation of the population dynamics of the two inoculated phages.

**Figure 1.**
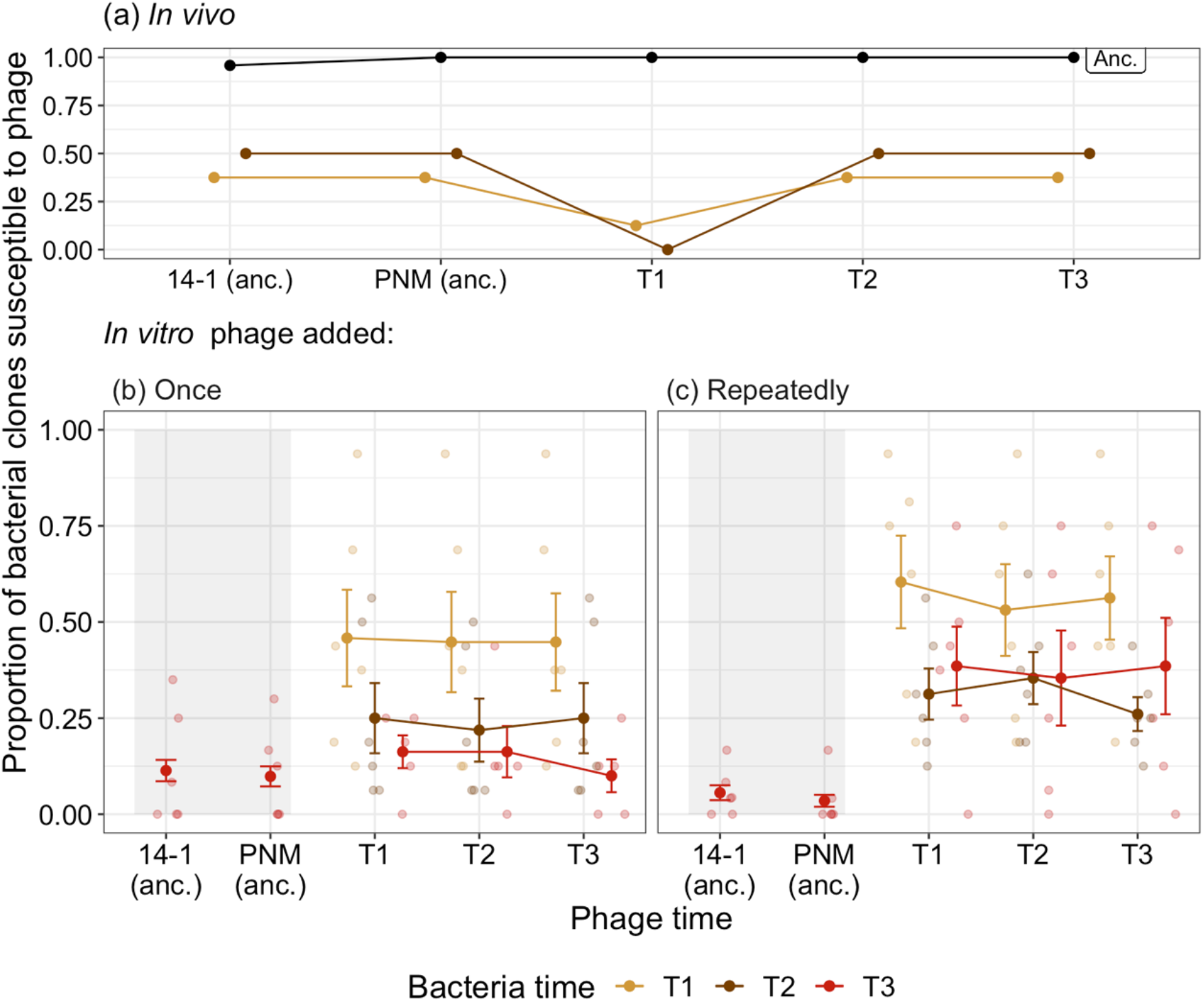
Time-shift assays of bacterial evolution and phage during phage therapy. The proportion of phage susceptible bacterial isolates from (a) ancestral (pre-phage therapy, “Anc.”), and days 2 (T1) and 4 (T2) of *in vivo* (clinical) phage therapy (n = 8) tested against ancestral phage (“anc.”: 14-1 and PNM) and phage isolates from days 2 (T1), 4 (T2) and 7 (T3). The same ancestral bacterial isolates are evolved in vitro with phage added either (b) once, at the start of the experiment for comparison to (c) in which phage are added repeatedly, emulating clinical treatment. For (b) and (c) the proportion of bacterial isolates389 susceptible to phage at three time-points (T1 - T3: days 4, 8 and 12 of coevolution) is presented with small 390 points as independent treatment replicates. Resistance of bacteria isolated at T3 to ancestral phage (“Anc.”: 391 14-1 and PNM) is indicated within the shaded region, note that these isolates are distinct from those tested against phage from T1-T3. Filled points are the mean proportion of resistant isolates across replicates. Bars = ±SE.

#### In vitro

We used the same bacterial isolates and phages to determine how well the *in vivo* evolutionary dynamics paralleled *in vitro* experiments in the lab. To assess the impact of phage application schedule, we added phages once at the start of the experiment, repeatedly (as was the case *in vivo*) or not at all. The dynamics in the phage treatments largely mirrored the *in vivo* dynamics. Resistance to phage increased through time, reaching comparable levels to that observed *in vivo* (>50%). While phage resistance increased in both phage treatments (ANOVA comparing models with and without bacterial time: phage added: once 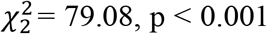 repeatedly 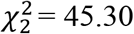, p < 0.001; Figure 1b, c), there was a significantly higher proportion of phage resistant bacteria in the treatment where phages were added once (90.81%, ±SE = 0.0003) compared to when phages were added repeatedly (66.7%, ±SE = 7.64; Tukey HSD comparing phage added once and repeatedly treatments: estimate = -1.63, z-ratio = -3.75, p = 0.0002; Figure 1b, c). All isolates from the control condition were susceptible to ancestral phages.

The lower resistance of bacteria when phages were added repeatedly was driven by evolution of increased phage infectivity. When phages were added once, resistance to ancestral phage (14-1: 88.64%, ±SE = 2.77; PNM: 90.15%, ±SE = 2.60) did not differ relative to phage isolated at the end of experimental evolution (90.0%, ±SE = 4.27; Tukey HSD comparing end-point phage resistance to contemporary phage to resistance to 14-1 (p = 1.0) and PNM (p = 0.938; Figure 1b). However, when phages were added repeatedly, resistance to ancestral phage (14-1: 94.41%, ±SE = 1.93; PNM: 96.5%, ±SE = 1.54) was significantly higher than to phage isolated at the end of experiment (61.5%, ±SE = 12.5), indicative of increased phage infectivity (Tukey HSD comparing end-point phage resistance to contemporary phage to resistance to 14-1 (p < 0.001) and PNM (p < 0.001); Figure 1c). We saw no further evidence of phage evolution in either treatments, with phage resistance being independent to the time point from which phage were isolated during experimental evolution (ANOVA comparing models with and without phage time: phage added: once 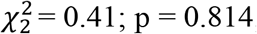; repeatedly 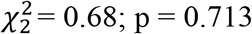; Figure 1c). This suggests the increase in phage infectivity evolved before the first sampling time point (the second transfer, or 96 hours). The absence of any further changes in phages during experimental evolution did not warrant further investigation of the population dynamics of the two inoculated phages.

### Phage resistance-virulence trade-offs

We next assessed whether phage application altered the virulence of *P. aeruginosa* populations *in vivo* and *in vitro* using a *G. mellonella* infection model. Here, virulence can be calculated as the inverse of *G. mellonella* survival over time: the more virulent the bacterial isolate, the shorter the time until death. *In vivo*, phage therapy reduced bacterial virulence by 95.5% on average (median hazard ratio = 0.045, 95%CI = 0.012 – 0.163). When inoculated with bacteria isolated before phage therapy, 479 (99.7%) of *G. mellonella* died within 46 hours, with death occurring at a mean time of 15 hours (Figure 2a). In contrast, after phage therapy, bacteria killed only 70% of *G. mellonella* within 50 hours, with mean time of death occurring at 20.5 hours (Figure 2a).

**Figure 2.**
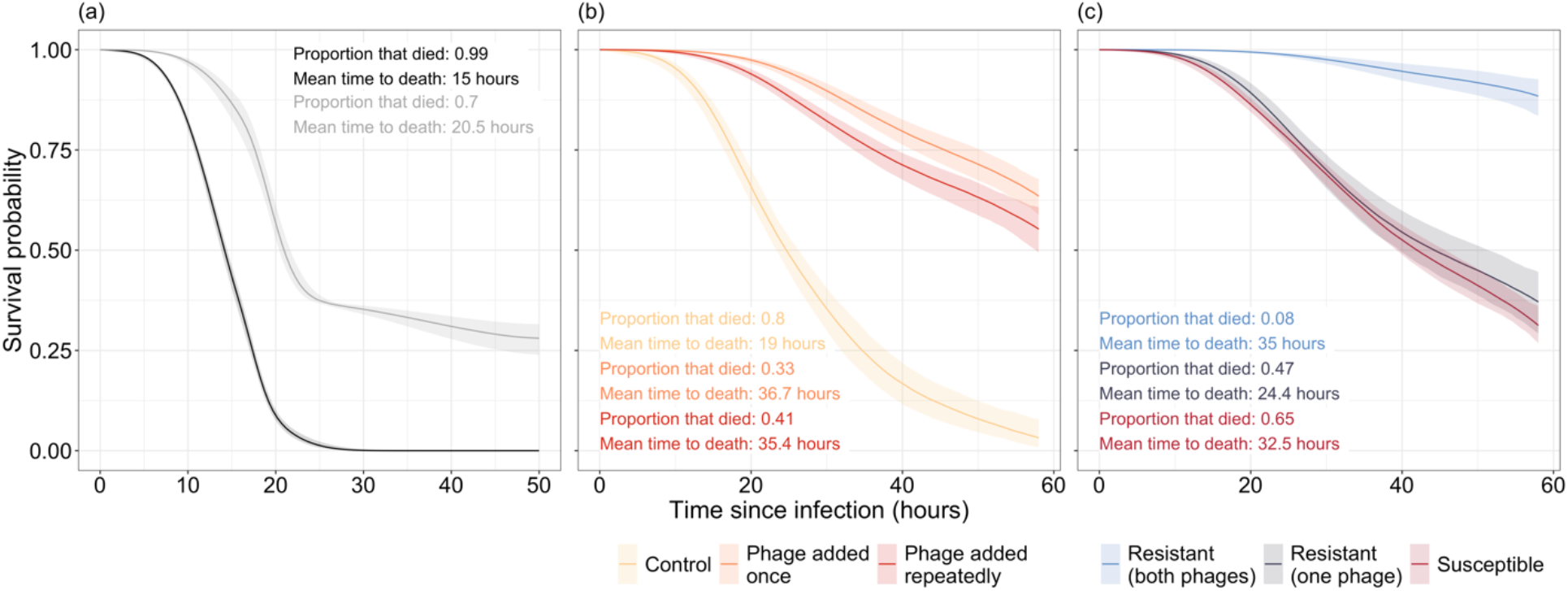
Survival curves of G. mellonella inoculated with P. aeruginosa isolates isolated (a) *in vivo* and (b, c) in vitro. (a) Virulence of bacterial isolates isolated from before (black) and after (grey) phage therapy. (b) Virulence of bacterial in control populations where no phage is present (yellow) is much greater than populations which were exposed to either a single (orange) or repeated (red) phage applications. (c)Virulence is significantly reduced in bacterial isolates resistant to both phages while bacteria resistant to one phage are non-significantly different in virulence to phage susceptible isolates. Lines represent the average prediction and shaded bands represent the 95% credible intervals of those predictions. Proportion and mean time to death of Galleria that died are presented as summary statistics for each treatment.

We also saw a reduction in virulence *in vitro* when populations had evolved with phage compared to without phage. Bacteria in the phage added once and phage added repeatedly treatments were 96.15% (median hazard ratio = 0.0385, 95%CI = 0.011 – 0.124) and 89.8% (median hazard ratio = 0.102, 95%CI = 0.0294 – 0.355) less virulent on average compared to the no-phage control populations (Figure 2b). In the control populations, 96 (80%) of the *G. mellonella* died within 47 hours, with death occurring at a mean time of 19 hours. In contrast, only 32.5% and 41.2% of *G. mellonella* died in the phage added once and phage added repeatedly treatments respectively, with a mean time to death of 36.7 and 35.4 hours (Figure 2b).

This difference was driven by two impacts of phage application. First, there was a huge reduction in virulence of bacteria that were resistant to both phages. When resistant to both phage, virulence reduced by 95.69% compared to susceptible bacteria (median hazard ratio = 0.0431, 95%CI = 0.0139-0.122) and was 93.33% lower than bacteria resistant to only a single phage (median hazard ratio = 0.0663, 95%CI = 0.0179-0.247; Figure 2c). Meanwhile, there was no reduction in virulence for bacteria resistant to just a single phage (median hazard ratio = 0.653, 95%CI = 0.185 -2.09) when compared to phage susceptible bacteria. Second, even susceptible bacteria became less virulent after phage application, with a 91.14% median hazard ratio = 0.0886, 95%CI = 0.0246-0.344) reduction in the phage added once treatment and an 80.7% reduction when phage was added repeatedly (median hazard ratio = 0.193, 95%CI = 0.04-1.06; Figure 2c).

### Growth rate

Evolving resistance against phage can trade-off with growth rates (Buckling et al., 2006; Koskella et al., 2011; Wright et al., 2018), and could be a plausible mechanism for virulence reduction. In the case of the *in vivo* isolates, the growth rate of pre-phage treatment isolates 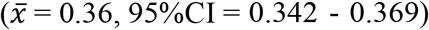 was significantly higher than that of post-treatment isolates 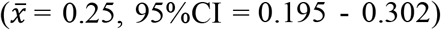 Post-treatment isolates which evolved resistance (to one or both phage) had a significantly lower growth rate 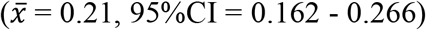 than the ancestral isolates 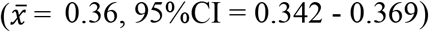 However, the 95% CIs of the susceptible isolates’ growth rate 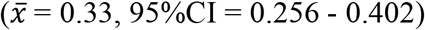 overlapped with the ancestral and resistant isolates, indicating non-significance, which was likely driven by a lack of statistical power (n = 3 susceptible isolates) (Figure 3a).

**Figure 3.**
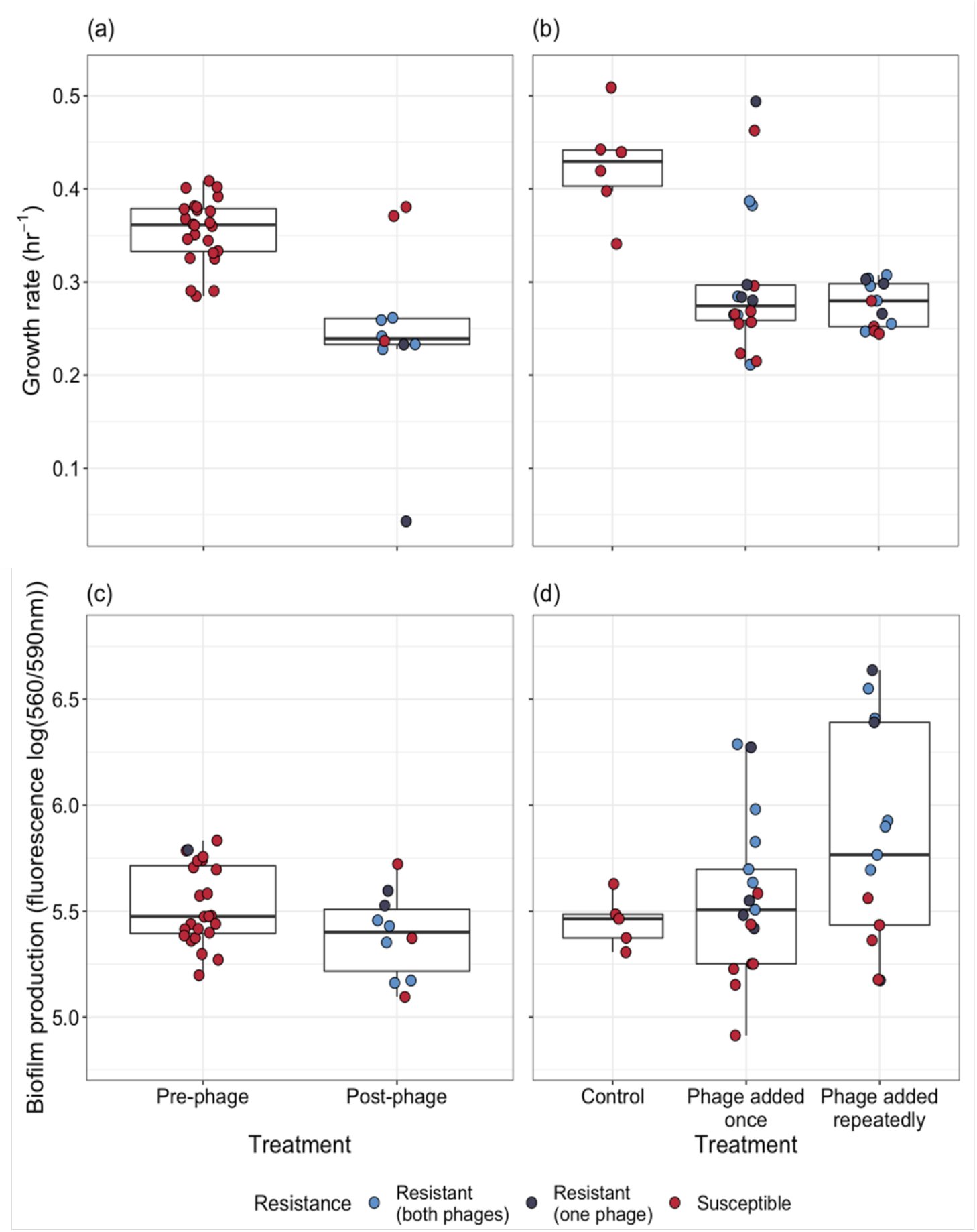
Phenotypic changes underpinning the resistance – virulence trade-off of bacterial populations. Populations were evolved (a, c) *in vivo*, and (b, d) in vitro. (a, b) Changes in bacterial growth were estimated for populations isolated (a) before and after phage treatment, and (b) between control and the phage treatment groups. Single points represent individual bacterial isolates of differing phage resistance levels: resistant to both phage (light blue), resistant to 14-1 or PNM (dark blue) or phage susceptible (red). Tops and bottoms of the bars represent the 75th and 25th percentiles of the data, the middle lines are the medians, and the whiskers extend from their respective hinge to the smallest or largest value no further than 1.5* interquartile range.

For *in vitro* isolates, growth rate was significantly affected by treatment 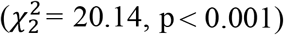 with the control group having significantly higher growth rates 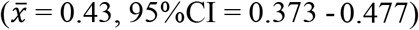 compared to treatments in which phage were added once 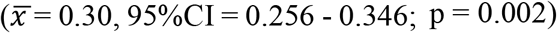 or repeatedly (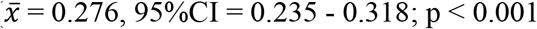; Figure 3b). No significant difference was observed in growth rates between each phage treatment (Tukey HSD: estimate = 0.025, t-ratio = 0.95, p = 0.628); and changes in growth were not associated with phage resistance (interaction between treatment and resistance: 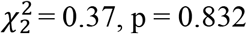; fixed effect of resistance: 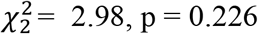; Figure 3b).

### Biofilm production

Biofilm production is often altered in phage resistant isolates (Harrison et al., 2015; Kim et al., 2015; Scanlan and Buckling, 2012) and this can also affect virulence and growth rate (Li et al., 2008; Wang et al., 2011). Therefore, we compared the biofilm productivity between isolates evolved *in vitro* and *in vivo*. For *in vivo* isolates, there was no significant difference in biofilm production between isolates isolated pre- 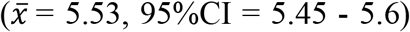; or post-phage therapy 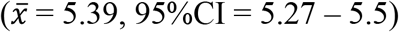, and between phage resistant 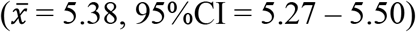 and susceptible isolates (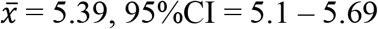 Figure 3c).

For *in vitro* isolates, there was no significant difference in biofilm production between treatments (treatment x resistance interaction: 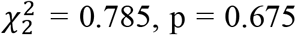; treatment: 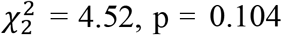). However, phage resistance was associated with increased biofilm production 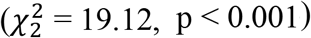, with phage susceptible isolates, 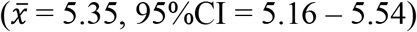 having lower biofilm production than isolates resistant to one 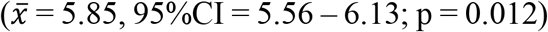 or both phages (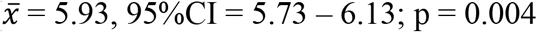; Figure 3d).

### Genetic basis of evolutionary changes

Given the parallel phenotypic differences we observed under *in vivo* and *in vitro* conditions, we next explored whether the underlying genetic changes were the same for *in vitro* and *in vivo* evolved populations. From the *in vitro* pooled sequencing, we found significant changes in 27 different genes with known function (Figure 4), 15 of which were changes in frequency compared to the ancestral population while 12 were novel mutations.

**Figure 4.**
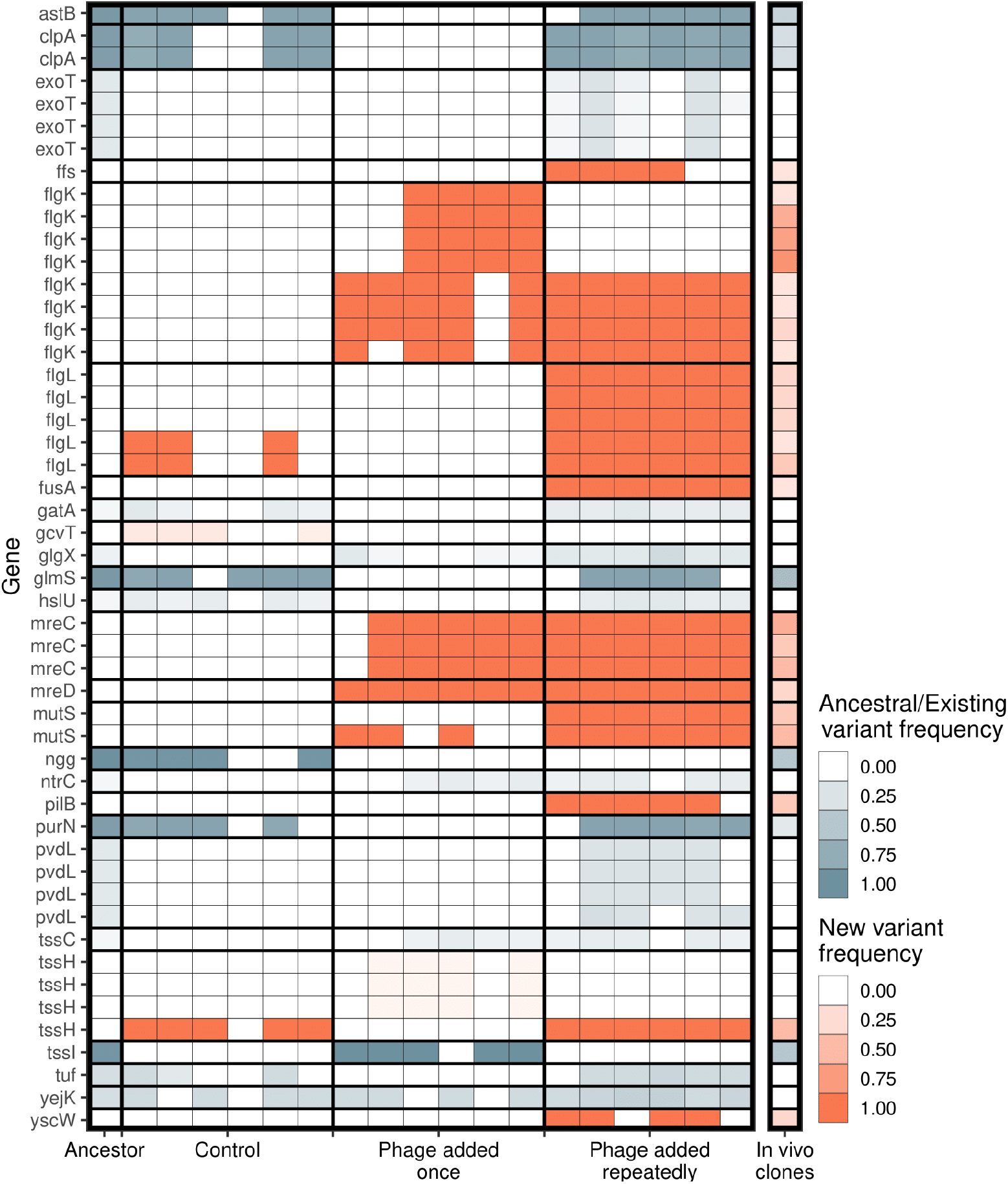
Significant genetic changes identified *in vitro* and *in vivo*. The frequency of genes that differed significantly between treatments *in vitro* are shown. Genetic changes that were already present in the ancestral population (first column) are shown in blue, while *de novo* mutations are in orange. Only genes whose function was identified are included. Multiple rows of the same gene indicate different genomic variants of the same gene. Colour intensity shows how frequent the genetic change is in the population. For *in vitro* treatments, each column is an independent replicate representing the results of the pooled sequencing. For the *in vivo* isolates (final column), all isolates were pooled to visualise the frequency of each genetic change *in vivo*.

In terms of genetic distance from the ancestral population, populations in the treatment where phage was added repeatedly diverged more than when phage was added once (Tukey HSD: *t* = -4.32, d.f. = 15, p = 0.0017), which diverged more from the ancestral populations than the control populations (Tukey HSD: *t* = -8.81, d.f. = 15, p < 0.0001; Figure 5a, b). Consistent with this, more novel SNPs/indels were found in the treatment populations where phages were added repeatedly (mean = 19.2) compared to where phages were added once (mean = 11.8, Tukey HSD: *t* = -4.80, d.f. = 15, p = 0.0006), which in turn had more novel SNPs/indels than control populations (mean = 2.5, Tukey HSD: *t* = -6.105, d.f. = 15, p = 0.0001; Figure 5b). Within-population diversity was highest in phage added repeatedly treatment (Tukey HSD phage added repeatedly *vs*. phage added once: *t* = -5.96, d.f. = 15, p = 0.0001; Tukey HSD phage added repeatedly *vs*. control: *t* = - 4.33, d.f. = 15, p = 0.0016), but there was no difference between control populations and phage added once treatment (Tukey HSD: *t* = -1.14, d.f. = 15, p = 0.2625; Figure 5c).

**Figure 5.**
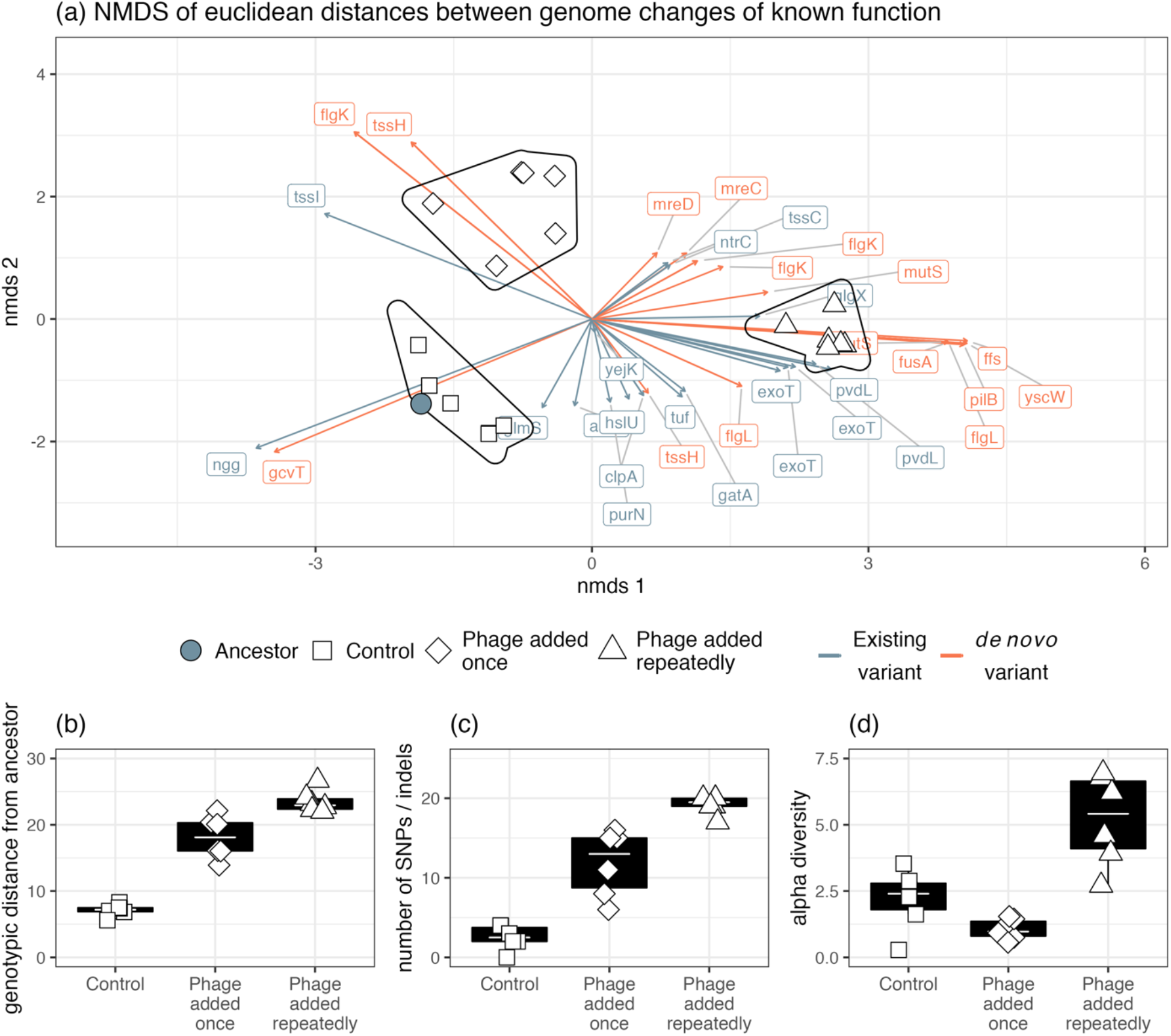
Genomic changes for populations evolved *in vitro*. (a) The divergence of each treatment group as indicated by Euclidean distances of each gene from the ancestral isolates. Genetic changes were characterised as a change in frequency of an existing gene in the ancestor or selection on a mutation that was gained during experimental evolution. The presence of phage significantly affected (b) genotypic distance from the ancestor, (c) the number of new SNPs and (d) alpha diversity.

*In vitro* treatment significantly altered genetic composition (i.e. the centroids of phage added once, phage added repeatedly and control populations were different, Figure 4, PERMANOVA, F_3,15_ = 16.05, R^2^ = 0.76, p = 0.0001). There was no difference in beta-diversity (calculated from distance-to-centroids between groups) between treatments (homogeneity of multi-variation dispersion ANOVA: F_3,15_ = 1.65, p = 0.22). Genetic changes associated with both *in vitro* phage treatments, but not control, were flagellar hook-associated protein (*flgK*) and rod-shape determining proteins (*mreC* and *mreD*) (Figure 4). Additionally, genetic changes in DNA mismatch repair (*mutS*), elongation factor G (*fusA*), signal recognition particle (*ffs*), type 3 secretion system pilotin (*yscW*), and pilus assembly protein (*pilB*) were mainly found *in vitro* in the phage added repeatedly treatment (Figure 4). We examined whether these same mutations arose in the clonal *in vivo* isolates. Of the 50 genetic changes identified *in vitro*, the *in vivo* isolates had a mean of 11.8 changes (maximum = 23, minimum = 1). Changes in frequencies or mutations were found at 16/27 of the same genes identified *in vitro* in at least one isolate (Figure 4). Crucially, some of the *de novo* mutations that were at high frequency in vitro were also at high frequency in vivo (*flgL* = 30%, *mutS* = 45%, *mreC* = 50%, *mreD* = 30%, *pilB* = 40%). We next assessed whether mutations arising in individual isolates *in vivo* could be linked to phenotypic changes (resistance, virulence, biofilm productivity, growth rate). If strong genotypic – phenotype links can be made, we would expect isolates to cluster according to their trait value. However, we found no obvious genotype to phenotype links *in vivo* and due to the small number of isolates, we were unable to conduct robust statistical analyses (Figure S2).

## Discussion

Highly parallel evolutionary dynamics were found *in vivo* and *in vitro*, showing that laboratory studies can be predictive of clinical phage therapy (phage-mediated decolonisation). Levels of phage resistance were comparable *in vivo* and *in vitro* with limited phage infectivity evolution occurring only *in vitro*. Additionally, phage resistance was associated with trade-offs with virulence and bacterial growth and similar mutations or genetic variants could explain these parallel evolutionary changes in both ecological contexts.

Parallel evolution is promising for future clinical work in which the trajectory of bacteria and phage (co)evolution may be predicted for individual patients. Given the difference in environments between that of a laboratory and human nasal cavity, it was surprising that evolution was so similar. The nasal cavity would have been more nutrient limited and spatially heterogenous and such conditions have been shown to change bacteria – phage interactions compared to evolution in laboratory growth medium. For instance, bacteria – phage coevolution is altered in soil compared to growth medium because of increased costs of resistance in the former (Gomez and Buckling, 2011); and in plant leaves, bacteria and phage have failed to evolve while otherwise coevolving in the laboratory due to lower bacteria – phage contact rates (Hernandez and Koskella, 2019).

Rapid phage resistance has been highlighted as a concern for phage therapy development (Torres-Barceló, 2018) and was found in this case study. Without ongoing reciprocal phage evolution, phage resistance is likely to lead to regrowth of bacteria during infections; this has been found in a few case studies but has been combatted by selecting a different phage (single or cocktail) (Aslam et al., 2020; Schooley et al., 2017; Wu et al., 2021; Zhvania et al., 2017). That resistance emerged predominantly by *de novo* mutations and not selection on existing variation further highlights the fact that stressful *in vivo* conditions need not prevent bacteria from rapidly adapting to phage. However, phage resistance was found to have growth rate costs which may further explain why some isolates remained sensitive to phage, and why decolonisation was ultimately successful. The natural nasal microbiota may have further contributed to this success by outcompeting slower growing phage resistant isolates (Wang et al., 2019). Such resistance – virulence trade-offs have been found in previous *in vitro*, animal model and clinical studies and is expected to improve treatment success in spite of phage resistance (Oechslin, 2018). Phage resistance however is not always costly and likely depends on the genetic background of the bacteria and phage, as well as the environmental context (Meaden et al., 2015; van Houte et al., 2016). The absence of clear links between resistance phenotypes and growth rate within our *in vitro* treatments, despite clear between-treatment effects, is presumably because the cultures were initiated with genetically diverse clinical isolates, rather than a single ancestral clone. Ultimately, phage resistance was beneficial in controlling the pathogen in this case study and such benefits could have led to treatment success.

Population genomics of the isolates *in vivo* and *in vitro* revealed parallel genomic changes and the effects of phage selection at a molecular level. In line with previous studies, phage selection accelerated molecular evolution which resulted in populations being more diverged from their ancestral pool than control isolates (*in vitro*) (Morgan et al., 2010; Pal et al., 2007). This suggests that phage placed strong selection on bacteria and this further resulted in mutator strains (*mutS* genotype), especially where phage were added repeatedly *in vitro* and *in vivo*, that likely accelerated the rate of evolution further (Morgan et al., 2010). Between both *in vivo* and *in vitro* results, mutations selected by phage gave insight into potential mechanisms of resistance. In particular, mutations to *mreD* and *mreC* encoded for cell shape (Cabeen and Jacobs-Wagner, 2005) while missense mutations at *flgK* and *flgL* probably resulted in flagella loss (Macnab, 2003). The result of these mutations was likely reduced cell surface area which reduces the probability of phage absorption (Dennehy and Abedon, 2021). *pilB* mutations were also selected which is likely a surface mutation to prevent PNM (a pilus binding phage) from attaching (Ceyssens et al., 2011).

Strong genotype – phenotype links were difficult to distinguish as a high number of the same mutations or existing variants were selected upon both *in vivo* and *in vitro*, and no unique candidate genes were obvious to explain phenotypic differences (i.e. growth rate, biofilm production). Flagella and *mreC* mutations are associated with increased biofilm production (Whiteley et al., 2001) and as they were found in the *in vitro* isolates, this may suggest a generalistic phage defence strategy (Harper et al., 2014). However, as these same mutations were found *in vivo*, without increases in biofilm production, strong genotype – phenotype links are unclear. Surface mutation is a common form of phage resistance (van Houte et al., 2016) but here it appears coupled with mechanisms that prevent absorption more generally; combined, these strategies could have reduced bacterial growth and/or virulence by impairing swarming motility (Haiko and Westerlund-Wikström, 2013). In contrast, other studies using the same phage but *P. aeruginosa* PA01 instead reported mutations predominantly at pilus and lipopolysaccharide receptors, showing that greater selection is placed on specific phage resistance strategies at phage binding sites (Wright et al., 2019, 2018). Our results highlight the importance of genetic background, and hence the use of clinical isolates over laboratory clones, in determining evolutionary trajectories (Betts et al., 2014; Friman et al., 2016; León and Bastías, 2015; Rossitto et al., 2018; van Houte et al., 2016).

How phages were applied *in vitro* had an impact on phage resistance. Although we observed a greater number of mutations where phages were added repeatedly (mimicking multiple phage doses), the proportion of resistant bacteria at the end of the experiment was significantly lower (61.5%) compared to where phages were just added at the start of the experiment (90%). That resistance of evolved bacteria to the ancestral phages (∼90%) did not differ between treatments suggests that differences in resistance to evolved phages was caused by phage infectivity evolution in the repeated phage treatment. This infectivity evolution occurred before the first time point, with no subsequent changes, and was presumably the result of greater mutation supply rate in this repeated phage application treatment. Multiple (Manohar et al., 2018) and higher (Barrow et al., 1998; Biswas et al., 2002; Debarbieux et al., 2010; Merril et al., 1996; Sunagar et al., 2010; Vinodkumar et al., 2008; Wang et al., 2006; Wills et al., 2005) phage doses have been shown previously to decrease morbidity and mortality in animal models. Our results build upon this work to show that maintaining high phage titres may have a positive effect in reducing resistant populations despite increasing the strength of selection for resistance.

Although our results suggest that results of laboratory studies may parallel phage resistance – virulence trade-offs during *in vivo* phage therapy, there a number of important caveats. Firstly, our *in vitro* – *in vivo* results were comparable with regards to the direction of change, but absolute effect sizes were greater *in vitro*. For instance, we observed a higher proportion of phage resistant isolates *in vitro* that were also far less virulent than isolates found *in vivo*. Therefore, it may be expected that phenotypic changes will be exaggerated *in vitro*. Secondly, as this patient had a nasal colonization, the role of the adaptive immune system would have been heavily reduced (Ooi et al., 2008). The immune system has the potential to improve phage therapy success by synergistically killing bacteria with phage which may diminish the potential for phage resistance by reducing bacterial growth (Leung and Weitz, 2017; Roach et al., 2017). Alternatively, immune cells may kill phage and therefore prevent phage acting on bacterial populations (Van Belleghem et al., 2018). Additionally, the nasal cavity has a diverse microbiome that may have contributed to *P. aeruginosa* decolonisation (Hu et al., 2018; Kumpitsch et al., 2019). The role of the microbiome will differ in infection type, with gut and respiratory infections potentially having a greater microbiome effect (Hu et al., 2018; Kumpitsch et al., 2019; Lloyd-Price et al., 2016) while septic infections may have a greater immune system effect (Van Belleghem et al., 2018). Therefore, it is important to compare laboratory vs *in vivo* phage therapy in a number of infection types. The genomic background of the bacteria may further affect how sensitive its resistance strategies are to the environment it is in as more costly resistance strategies are more likely to be selected *in vitro* than *in vivo* (van Houte et al., 2016). This encompasses not only genetic variation within species but also differences between species that may differ in infection sites, immune responses, virulence and sensitivity to phage (Abedon, 2019; Doffkay et al., 2015; van Houte et al., 2016). Further work using isolates from case studies and clinical trials will give an insight into the predictability of bacteria and phage evolution from laboratory to *in vivo* studies.

## Acknowledgements

Genome sequencing was provided by MicrobesNG (http://www.microbesng.com)

## Supplementary material

### Parallel resistance evolution to phages during clinical phage therapy and in vitro

#### Supplementary figures

Data cleaning of the growth rate assays. As shown in Figure S1, we observed a sharp increase in OD_600_ reads within the initial 2.5 hours which was separate from the exponential growth phase. To prevent the rolling regression taking these initial measurements into account, these values were removed from analyses.

**Figure S1.**
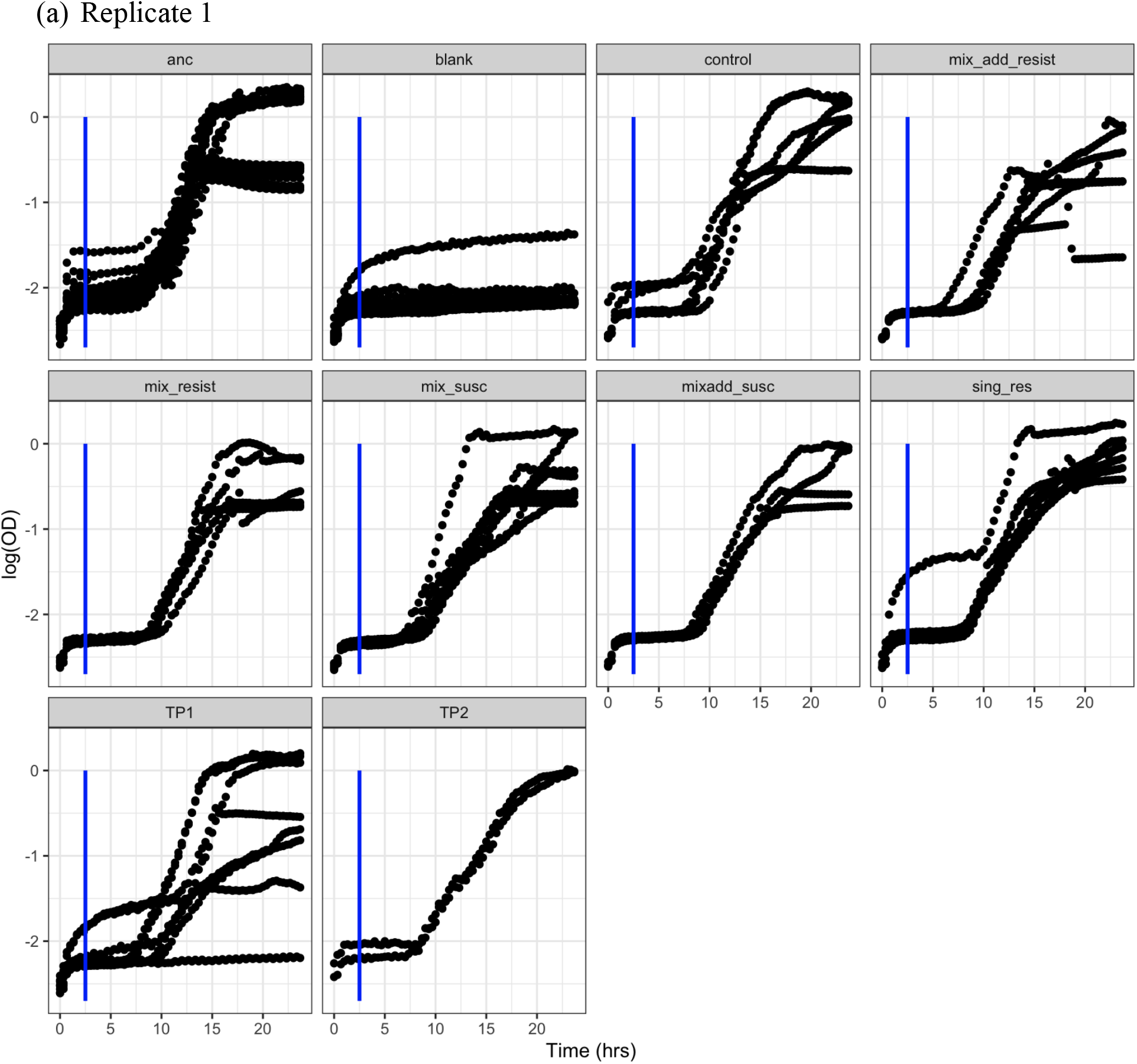
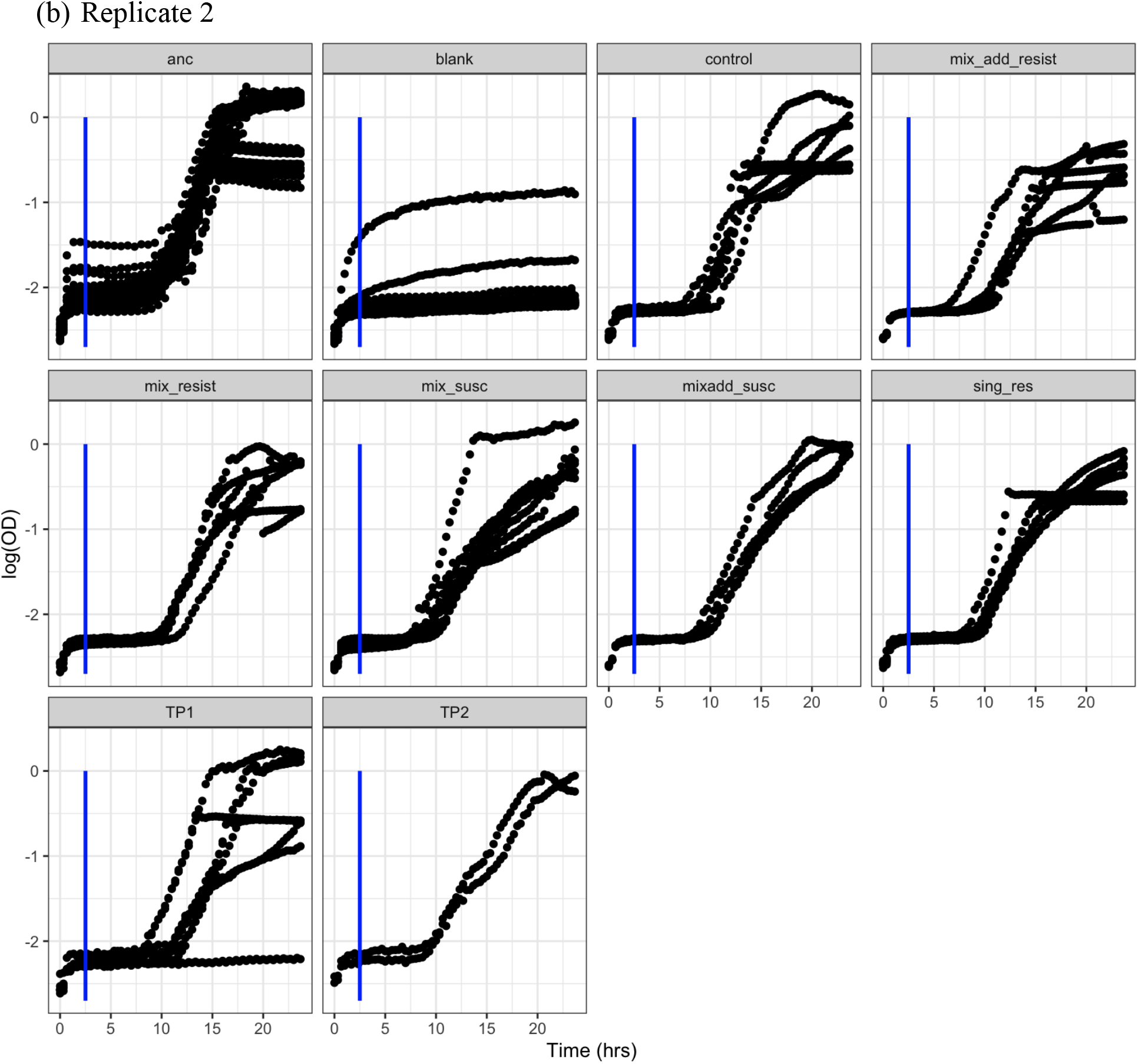
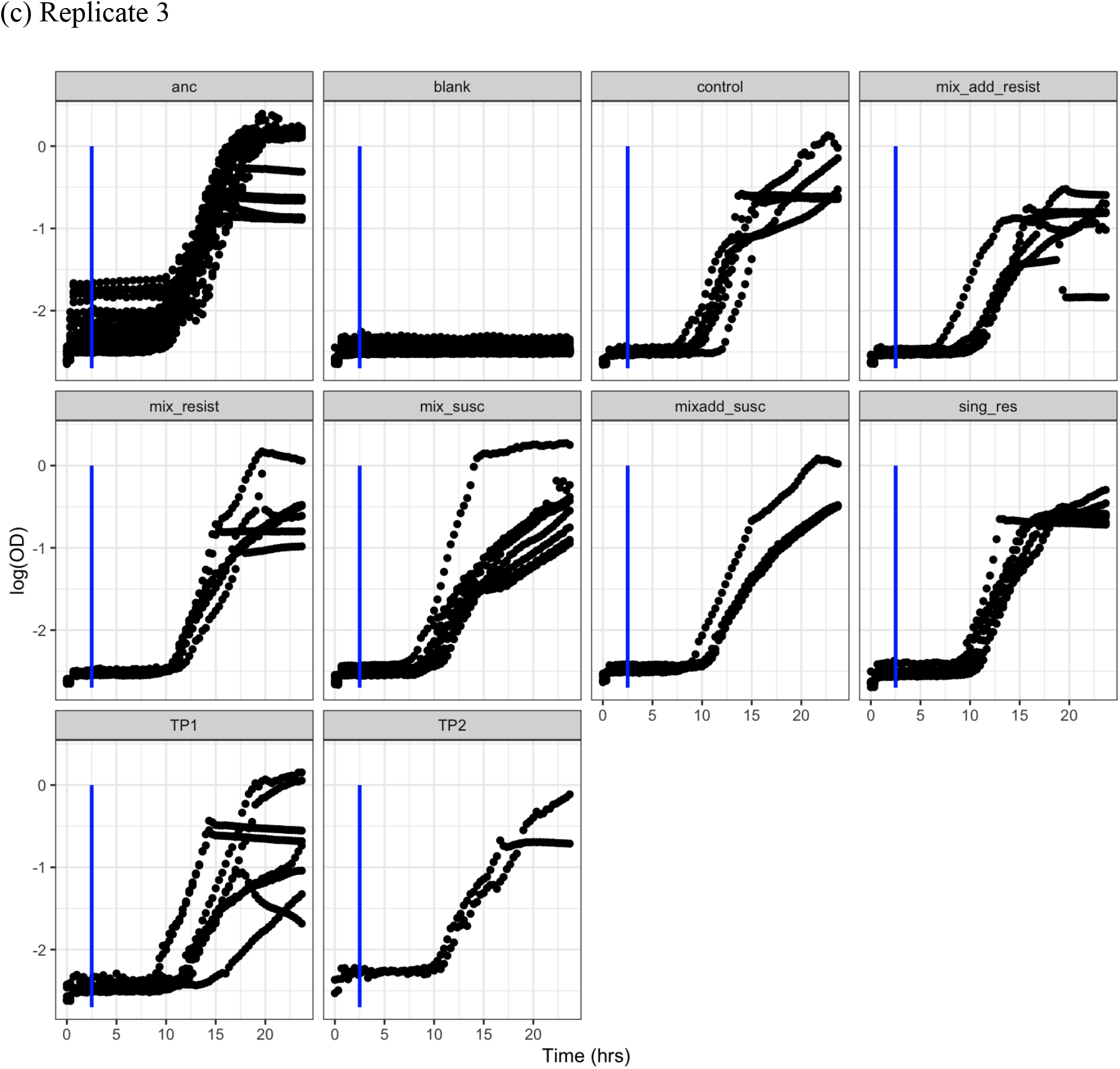
Growth curves of bacterial clones as measured by change in log-transformed ocular density (OD_600_) over time (hours). Measures from the first 2.5 hours were excluded (blue line) as a rise in OD was observed independent of bacterial logistic growth which may skew the rolling regression process. Growth curves were independently replicated a total of 3 times (a - c) owing to cellular clumping which may have skewed OD measures. (“anc” = ancestral clones; “blank” = uninoculated wells; “control” = control clones; mix add” = clones from repeated phage treatment; “mix” = clones from treatment where phage were added once; “resist” = phage resistant clones; “susc” = susceptible clones; “sing_res” = in vitro clones resistant to one phage; “TP1” = *in vivo* clones from day 2; “TP2” = *in vivo* clones from day 4).

**Figure S2.**
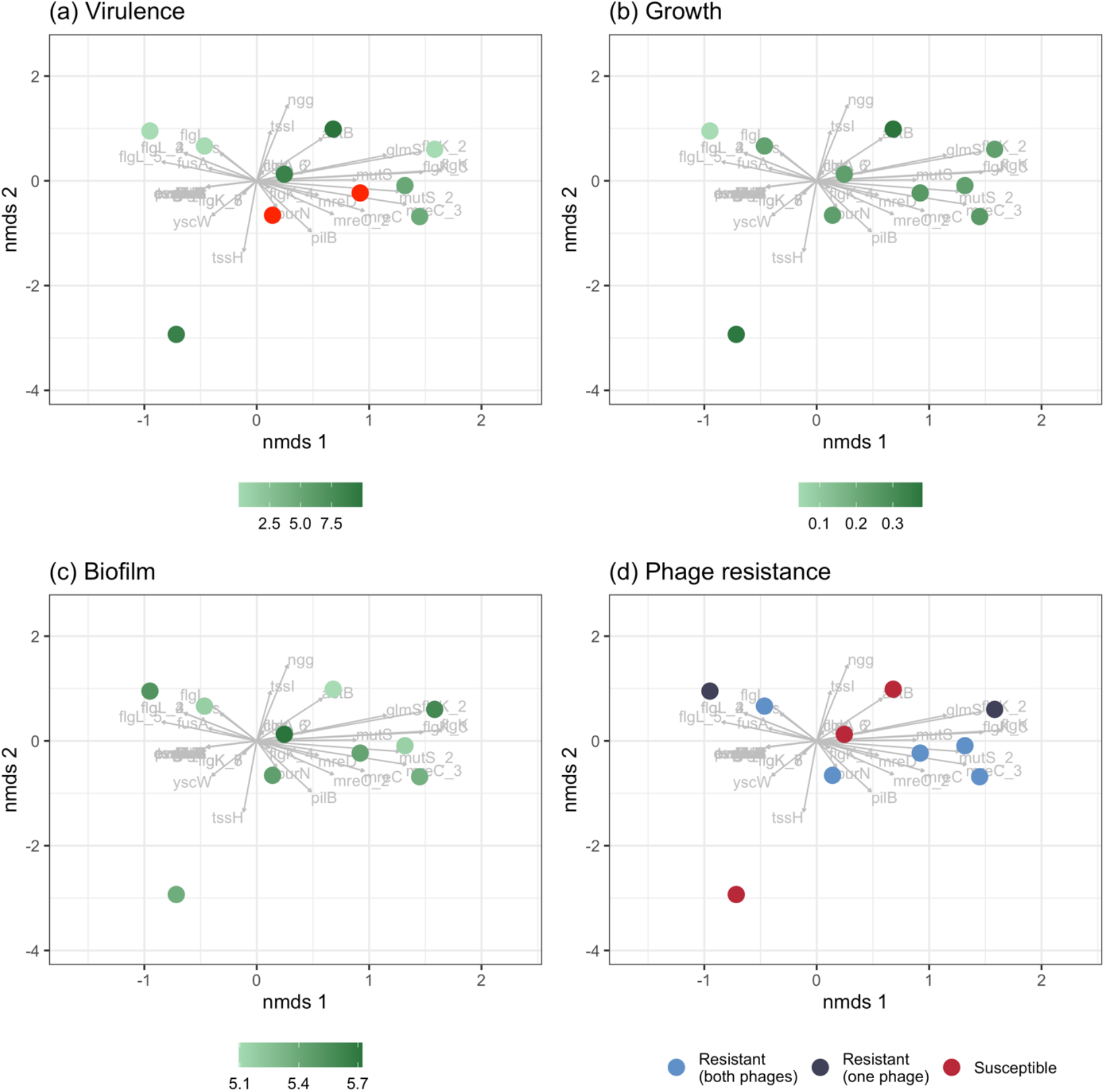
Associations between genetic similarity and phenotypic traits for *in vivo* clones. Individual points represent individual clones isolated at day 2 (n = 8) and day 4 (n = 2) of phage treatment. Genes that diverged in frequency *in vitro* from the ancestor are indicated in grey while *in vivo* clones are overlayed according to their individual genetic make-up. Across quantitative (a – c) and qualitative (d) phenotypic traits, no clustering in genotypes and clones of particular phenotypes is observed.

#### Supplementary Tables

**Table S1.** Susceptible colonies picked from *in vitro* phage treatments for phenotypic assays and sequencing.

| Phage resistance | Phage added | Treatment replicate | Clone number |
| --- | --- | --- | --- |
| Susceptible | Once | 6 | 1 |
| Susceptible | Once | 6 | 2 |
| Susceptible | Once | 6 | 3 |
| Susceptible | Once | 3 | 4 |
| Susceptible | Once | 3 | 5 |
| Susceptible | Once | 3 | 6 |
| Susceptible | Once | 5 | 7 |
| Susceptible | Once | 5 | 8 |
| Susceptible | Repeatedly | 6 | 9 |
| Susceptible | Repeatedly | 6 | 10 |
| Susceptible | Repeatedly | 6 | 11 |
| Susceptible | Repeatedly | 1 | 12 |

**Table S2.** Pairwise comparisons of resistance of *in vivo* clones from day 2 against phage from contemporary (day 2) and two future time-points (days 4 and 7). P-values adjusted using the Tukey method of comparing a family of three estimates. “SE” = standard error.

| Contrast | Estimate | SE | z-ratio | p-value |
| --- | --- | --- | --- | --- |
| Day 2 - Day 4 | - 16.5 | 10 | -1.65 | 0.2262 |
| Day 2 - Day 7 | - 16.5 | 10 | -1.65 | 0.2264 |
| Day 4 - Day 7 | 0.000001 | 5.32 | 0 | 1 |

## Supplementary methods

### Data cleaning of biofilm dataset

In the *in vitro* dataset, one replicate in the control and one in the susceptible treatments were found to be outliers that significantly impacted data analysis. In the susceptible treatment, the outlier (711) had a value 2.9x greater than the group mean (246.9) and this was 2.7x greater than the next nearest value (266.1). Similarly, in the control group, the outlier (708.6) had a value 2.26x greater than the group mean (313.5) and this was 2.5x greater than the next nearest value (278). We subsequently determined the impact of these values individually and when together on model output by sequentially removing these from the dataset (Table S3, S4). Resistance was significant regardless of outlier removal (Table S3) as were comparisons between resistant and susceptible groups (Table S4). Although treatment became significant when the susceptible outlier was removed, all Tukey-HSD comparisons were non-significant. The only change detected in Tukey HSD comparisons was that between the single resistant and susceptible isolates which was found when the susceptible and both outliers were removed (Table S4).

**Table S3.**
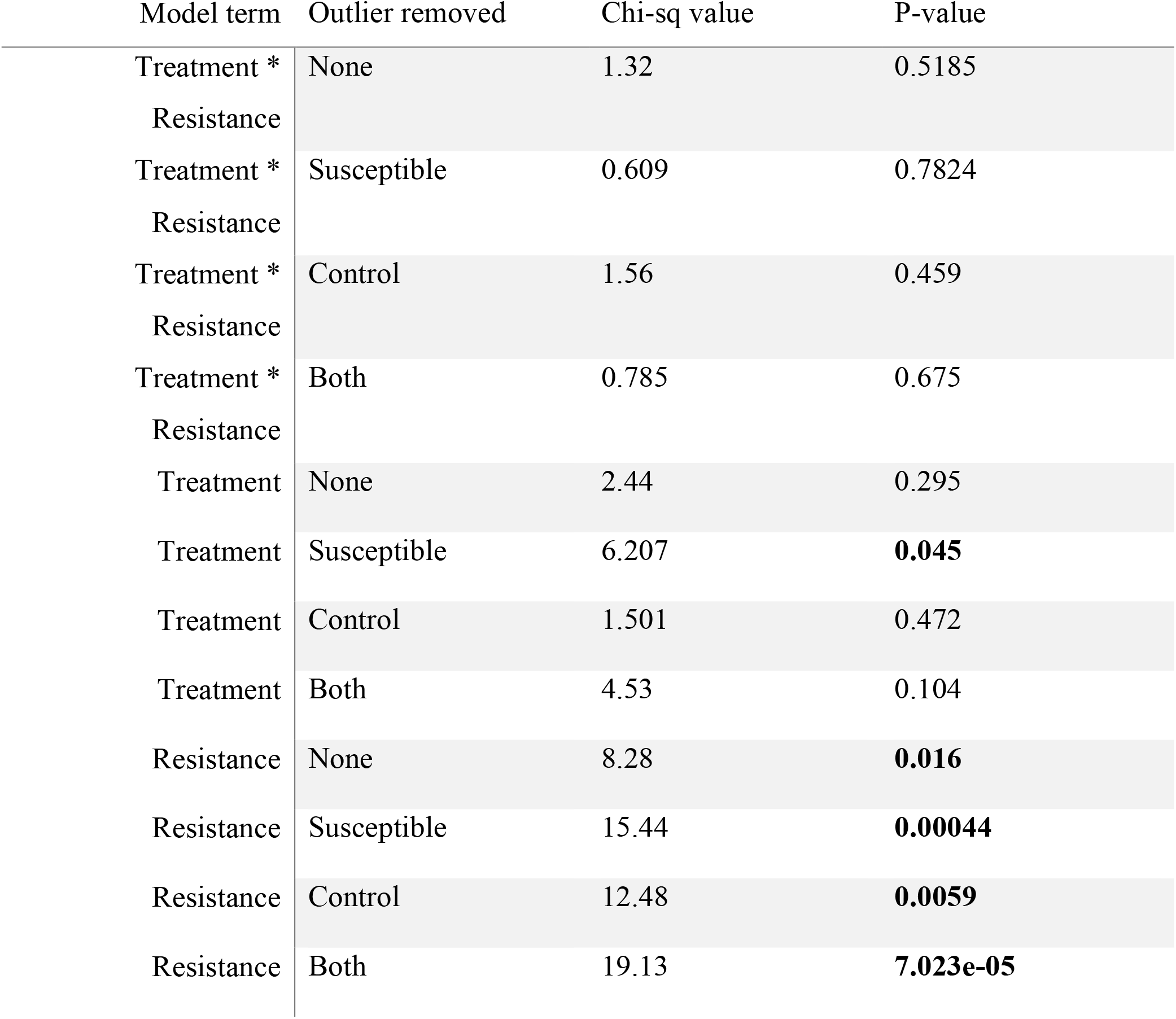
The effect of outlier removal on the significance of model terms (treatment and phage resistance). Outlier removed states in which treatment each outlier was identified and subsequently removed. Significant p-values in bold.

**Table S4.**
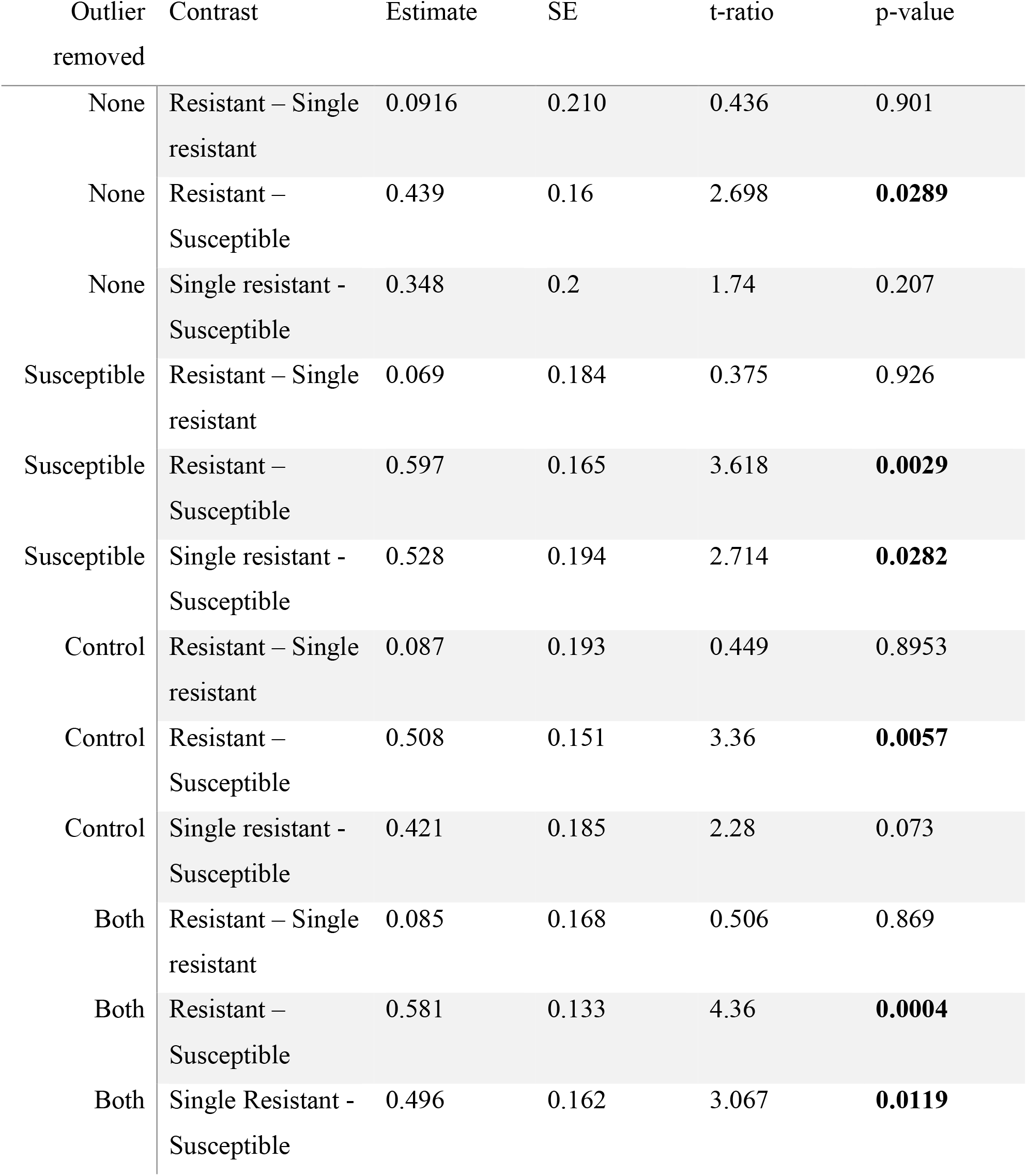
The effect of outlier removal on Tukey HSD contrasts comparing biofilm productivity between phage resistance groups. Significant p-values highlighted in bold.

| Outlier removed | Contrast | Estimate | SE | t-ratio | p-value |
| --- | --- | --- | --- | --- | --- |
| None | Resistant – Single resistant | 0.0916 | 0.210 | 0.436 | 0.901 |
| None | Resistant – Susceptible | 0.439 | 0.16 | 2.698 | **0.0289** |
| None | Single resistant - Susceptible | 0.348 | 0.2 | 1.74 | 0.207 |
| Susceptible | Resistant – Single resistant | 0.069 | 0.184 | 0.375 | 0.926 |
| Susceptible | Resistant – Susceptible | 0.597 | 0.165 | 3.618 | **0.0029** |
| Susceptible | Single resistant - Susceptible | 0.528 | 0.194 | 2.714 | **0.0282** |
| Control | Resistant – Single resistant | 0.087 | 0.193 | 0.449 | 0.8953 |
| Control | Resistant – Susceptible | 0.508 | 0.151 | 3.36 | **0.0057** |
| Control | Single resistant - Susceptible | 0.421 | 0.185 | 2.28 | 0.073 |
| Both | Resistant – Single resistant | 0.085 | 0.168 | 0.506 | 0.869 |
| Both | Resistant – Susceptible | 0.581 | 0.133 | 4.36 | **0.0004** |
| Both | Single Resistant - Susceptible | 0.496 | 0.162 | 3.067 | **0.0119** |

